# Suppression of fibrillation consisting of stable rotors by periodic pacing

**DOI:** 10.1101/2021.08.25.457679

**Authors:** Pavel Buran, Thomas Niedermayer, Markus Bär

## Abstract

Recent experimental studies have shown that a sequence of low-energy electrical far-field pulses is able to terminate fibrillation with substantially lower per-pulse energy than a single high-energy electric shock (see S. Luther et al. **Nature** 475 (7355), 235-239). During this low-energy antifibrillation pacing (LEAP) procedure only tissue near sufficiently large conduction heterogeneities, such as large coronary arteries, is activated. In order to understand the mechanism behind LEAP, We have carried out a statistical study of resetting a medium filled by one or more stable spirals (“rotors”) in a two-dimensional electrophysiological model of cardiac tissue perforated by blood vessels to the resting state (“defibrillation”). We found the highest success probabilities for this defibrillation for underdrive pacing with periods 10 – 20 percent larger than the dominant period of the stable rotors in the unperturbed dynamics. If a sufficiently large number pulses is applied and an optimal pacing period chosen, the energy per pulse required for successful defibrillation is about 75 - 80 percent lower than the energy needed for single-shock defibrillation. Optimal conditions to control and suppress fibrillation based on stable rotors, hence, are similar to the ones in found for the case of an electrophysiological model displaying spatiotemporal chaos (“electrical turbulence”) in an earlier study (see P. Buran et al. **Chaos** 27, 113110 (2017)). The optimal pacing period is found to increase with increasing strength of the electrical field strength used in the model. The success probability also increases strongly until the fourth or fifth pulse administered, which is strongly correlated to an observed increase of the fraction of re-excitable tissue with each subsequent pulse. Monitoring the fraction of excitable tissue in the model as key quantity of the excitable medium, moreover, enabled us to successfully predict the optimal pacing period for defibrillation.

## I. INTRODUCTION

A loss of rhythm and synchronization of the cardiac electrical activity, orchestrating heart contraction and the pumping of blood, is associated with a number of arrhythmias including atrial fibrillation (AF), ventricular tachycardia (VT) and ventricular fibrillation (VF). VF is a particularly dangerous malfunction of the heart, preventing an effective mechanical contraction of the ventricles and causing sudden cardiac death within a few minutes if left untreated. The electrical activity during VF is dominated by multiple stable spirals (rotor) or by multiple wavelets in a spatiotemporally chaotic state.^1–4^ VT on the other hand is often assumed to be generated by one single rotor; VT is not directly lethal, it can, however, degenerate rapidly to VF.

Defibrillation by a strong electrical pulse is the standard way of terminating VF. However, strong shocks are associated with adverse side effects including pain and trauma^5^ as well as damage of the myocardium^6^. These adverse effects could be potentially avoided or diminished if VF would be terminated reliably by alternative method using significantly lower energy. Recent experimental studies^7,8^ of AF *in vitro and in vivo* and VF *in vitro* have shown that a sequence of five or more farfield pulses with stimulation rates close to the arrhythmia cycle length typically require 80% − 90% less energy per pulse than needed for defibrillation with a single shock. This method has been named low-energy antifibrillation pacing (LEAP). A similar energy reduction was also found with much faster pacing rates than the cycle length for AF^9,10^, while the energy may be even reduced further by an order of magnitude for VT^11,12^.

Defibrillation is based on the fact that an electric field depolarizes and hyperpolarizes the tissue close to conductivity heterogeneities^13^ which act as virtual electrodes^14^. The strength of this effect depends both on the strength of the electrical field and on the size and shape of the heterogeneities.^15–17^ Only tissue at mayor conduction heterogeneities may be activated by low-energy pulses. Therefore, global tissue activation and wave termination often originate only from a few localized activation sites (hot spots). Luther *et al*.^8^ suggested that these hot spots are given by large coronary arteries and showed that the size distribution of the heterogeneities follows a power law. The results Caldwell *et al*.^18^, however, did contest such a co-localization of hot spots and major coronary vessels.

The mechanism how LEAP terminate arrhythmias is not well understood yet since experimental methods to visualize the whole three dimensional electrical activity inside the heart are missing. A new promising method which might give deeper insights in the future is the determination of the three dimensional electrical activity over a ultrasound measurement of the mechanical activity of the heart.^19^ Current theoretical approaches to understand LEAP often focus on the process of unpinning and removal of a small number of stable spirals.^15,20–22^ In these papers, it was demonstrated that spirals that were pinned at larger heterogeneities are best terminated if the LEAP pacing frequency is set to 80% − 90% of the rotation frequency of the spiral around the pinning site. This choice allows for an efficient scanning of the phase of the spiral in order to find the vulnerable window for spiral termination.^23,24^ Another common interpretation of the mechanism behind LEAP is synchronization, suggested by the observations in experiments^8,25^ and simulations^25,26^ wherein each pulse gradually entrains during successful LEAP more tissue until the entire tissue is synchronized and no excitation waves are left. In this studies LEAP was successful for overdrive^8,26^ as well as for underdrive^8,25^ pacing.

In a previous paper^27^ we have carried out a statistical study of success probabilities for low energy periodic pacing applied to states of spatiotemporal chaos within exemplary two dimensional electrophysiological models, in which the non-conducting heterogeneities were represented by circles whose size distribution was given by the experimental distribution of radii of coronary arteries reported in Ref.^8^. Two alternative cell models^28,29^ exhibiting similar length scales of electrical turbulence and the correct magnitude of the electric field necessary single pulse defibrillation were studied in detail. In both cases, a near-to-resonant pacing with the dominant period of unperturbed electrical turbulence results in maximal energy reduction already at moderate field strength. For a model featuring a pronounced dominant period, the reduction reached up to more than 98% per pulse compared to single pulse defibrillation. For larger field strengths, moderate underdrive pacing was found to be somewhat more efficient than resonant pacing.

In the present study we use again a simple two dimensional model of cardiac tissue perforated by blood vessels, however, focus on the termination of a regime of stable rotors, using a modified version of the Luo-Rudy model^28^ with parameter values from^30^. Therein, the defibrillation success probability of a periodic pacing protocol analogous to the LEAP method was determined starting from a state with a single free spiral, a single pinned spiral or multiple free spirals. The aim of this study is to get a deeper understanding of LEAP for stable spirals and to find the optimal protocol in this case. Therefore we have investigated the defibrillation success rate and other aspects of LEAP. It turned out that the optimal pacing period with respect to the termination probability is typically slighlty larger than the dominant period of the stable spirals (=rotation period), i. e. a mild so-called underdrive pacing provides optimal conditions to terminate the activity starting from a state of one or more stable rotating spirals. In addition, a high fraction of excitable tissue right before the application of the pulses was found to be reliable indicator for the success of defibrillation. This knowledge allowed for a computationally much more efficient alternative to estimate the optimal pacing period for LEAP.

In Sec. II, we introduce the the cellular model along with the numerical methods, our choice of initial conditions and for the periodic protocol. Sec. III contains the main results regarding single pulse high-energy defibrillation as a reference and defibrillation by low-energy periodic pacing as the main subject of this study. Sec. III concludes with the demonstration that the success of defibrillation by periodic pacing is strongly correlated with the amount of availabe excitable tissue, which depends on the time since the previous pulse and the number of pulses applied. We show that this observation allows a reliable prediction of the optimal pacing period with the highest success rate. Section IV provide a discussion of the results and a short conclusion, respectively.

## II. METHODS

We have performed extensive numerical simulations for defibrillation protocols with different electrical field strengths, number of pulses and pacing periods for states with multiple free stable spirals, one free stable spiral and one pinned stable spiral. In order to obtain statistically valid results, typically 50 simulation runs with different initial conditions were performed for each configuration. Therefore, we restricted the simulations to the same simple two-dimensional model of homogeneous, isotropic cardiac tissue, used in our previous work^27^, in which the heterogeneities are represented by circles whose size distribution is given by the distribution of radii of coronary arteries measured in Ref.^8^ and which bases on the monodomain model^31^ in conjunction with a cellular model.

### A. Numerical methods and model parameters

Simulations were carried out on a quadratic 2-dimensional equidistant finite-difference grid with no-flux boundary conditions and 1000 × 1000 nodes, resulting in a grid spacing of Δ*x* = 10^−3^ *L*, where *L* is the system size (side length of the quadratic tissue domain). Conductivity heterogeneities were included into this grid as non-conducting patches by modified Neumann-boundary-conditions^15,17^ and the time integrations were performed with a time step of 0.1 ms, which was sufficient to guarantee numerical stability.

In order to keep the results comparable with the ones in our previous work^27^, we used the exact same configuration of the simulation domain regarding system size *L* and distribution of heterogeneities. The system size of the quadratic tissue domain was *L* = 20 cm, which has then the extension of the order of magnitude of the surface of a (large) mammalian heart. The distribution of the heterogeneities followed the experimental size distribution *p*(*R*) ∝ *R*^−2.75^ of large coronary arteries given in Luther *et al*.^8^. As the cutoffs of the power law and as the density of all heterogeneities *R*_min_ = 3 × 10^−4^*L, R*_max_ = 4 × 10^−3^*L* and *ρ*_0_ = 1.6 × 10^4^*/L*^2^ were chosen. Heterogeneities with a diameter smaller than the grid spacing (2*R <* Δ*x*) where set to the size of one pixel. Note, that the area covered by defects amounts to around 3 percent of the total area used in the simulations in this paper. We have checked that different realization of positions and radii resulted in almost identical defibrillation successes. Therefore, the same realization of positions and radii was used in simulations while the initial states were varied unless stated otherwise.

As the cellular electrophysiological model, we used the Luo-Rudy model^28^ with parameter values from^30^, which exhibit stable spirals. This allowed us to study initial conditions with multiple spirals (associated with fibrillation) and single spirals either free or pinned (which are often associated with tachycardia), see Fig. 1. The diffusion constant with a value of *D* = 5.625 × 10^−3^*L*^2^/s was chosen such, that it resulted in a single defibrillation threshold that is consistent with experiments^8,32^, see Fig 3.

**FIG. 1.**
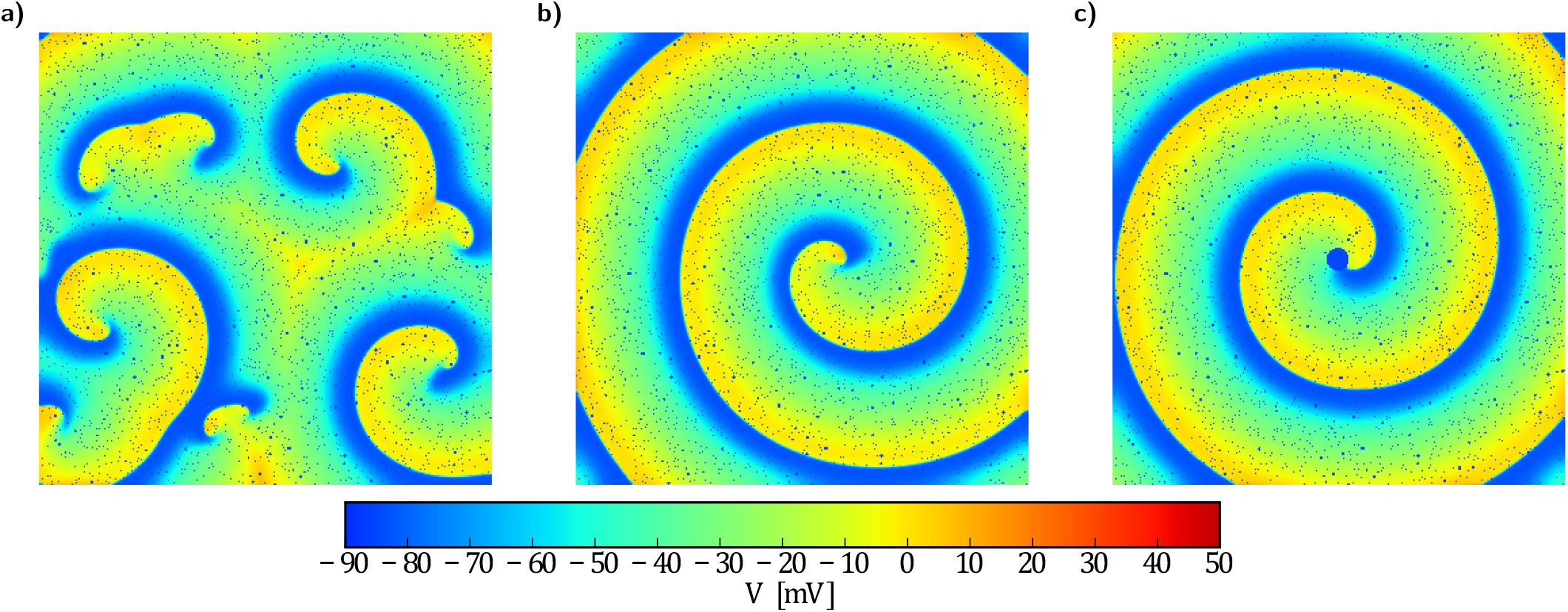
(a) Snapshot of transmembrane potential *V* for a state with multiple spirals, where an average of 10 phase defects are found. (b) Snapshot for a state with a single spiral, whose cores is located at the center. (c) Snapshot for a state with a single pinned spiral, whose cores is pinned to a circular heterogeneity at the center with a radius of 0.025 *L*. Conductivity heterogeneities are represented by small blue circles. Movies of the simulations are provided in the supplementary material.

### B. Determination of fraction of excitable tissue

The fraction of excitable tissue of a state was calculated as the number of excitable points divided by the total number of points. Excitable points were characterized by having a transmembrane potential lower than the activation threshold (*V*_thresh_ = 40 − mV) and releasing an action potential if the transmembrane potential exceeds this activation threshold. Therefore, additional time integration had been performed, where the transmembrane potential was set to the value of the activation threshold, and it was checked whether the transmembrane potential would achieve a value greater than 0.0 mV.

### C. Defibrillation protocols

State-of-the-art defibrillators execute a biphasic, asymmetric protocol.^33^ Nevertheless, we initially tested both monophasic and biphasic defibrillation protocols. For monophasic pulses, a rectangular waveform with a amplitude *E* and duration of 10 ms was chosen. The waveform for biphasic pulses consisted of two subsequent rectangular parts with identical amplitudes *E* and opposite field direction. We fixed the total pulse duration to 10 ms and found that 7 ms (forward) and 3 ms (backward) lead to the best defibrillation results, similar to our results results found in electrophysiological models exhibitng spatiotemporal chaos.^27^ The used protocols contained *n* identical biphasic pulses with a constant interval and the pacing period *T* is defined as the time difference between the onset of subsequent pulses.

### D. Generation of initial conditions

A single free spiral wave was initiated using the standard cross-field stimulation technique^34^. We run the simulation for 5 s to allow the spiral(s) to stabilize. Subsequently, we let the simulation run for one more complete rotation of the spiral to obtain the single spiral initial states at different equidistant phases. For the pinned spiral states, we placed one circular heterogeneity with a radius of 0.025*L* at the center of a single spiral state. Then, the simulation was carried out for 5 more s to allow the spiral to pin and to stabilize. Subsequently, we ran again the simulation for one more complete rotation of the pinned spiral to obtain the pinned spiral initial states at different equidistant phases. In order to get the multiple spirals initial states, we first cut from five randomly chosen single spiral states a quadratic section. Each section had a side length of 500 nodes (half of the system size) and its center was located in the middle of the single spiral, randomly additionally shifted up to 25 nodes in each direction. Then we placed in every quarter of a quiescent state one and at a random position the fifth of these five sections of cut spirals. Finally we run this generated states for 1 s to stabilize. In all cases, we generated 50 different initial states.

## III. RESULTS

### A. Spectra

Fig 2 shows the spectra for states with multiple spirals, a single spiral and a single pinned spiral. All three spectra are nearly identical and do not seem to depend on the number of spirals nor on the fact whether a spiral is pinned or not. They are all three characterized by a very narrow peak at *f*_0_ = 10 Hz or *τ*_0_ = 100 ms respectively corresponding to the rotation period of the spirals and further local maxima are at multiples of this frequency. The characteristic length scale of the unperturbed patterns is given by the wavelength of the spiral(s) that amounts to around 5.5 cm. This scale is almost two orders of magnitude bigger than the radius of the largest defect which equals 0.08 cm. As a result the defects in the medium do not change propagation properties substantially.

**FIG. 2.**
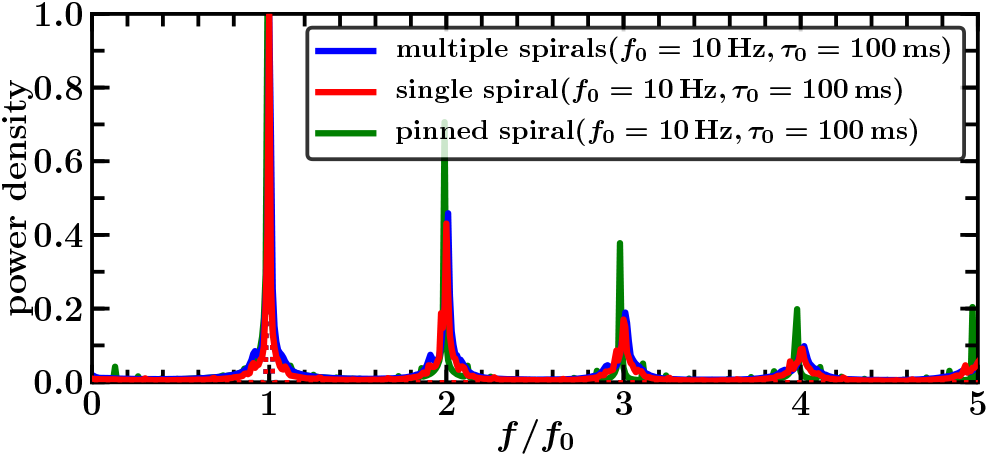
Scaled spectra of transmembrane potential *V* for multiple spirals (blue), a single spiral (red) and a single pinned spiral (green). Simulations were performed for 10 s and with 10 different initial conditions. The displayed graphs are averages of spectra of the transmembrane potential *V* at all spatial grid points and from all the 10 simulations runs. The spectra are in all threes cases nearly identical and characterized by a narrow peak at *f*_0_ = 10 Hz and further local maxima at multiples of this frequency.

**FIG. 3.**
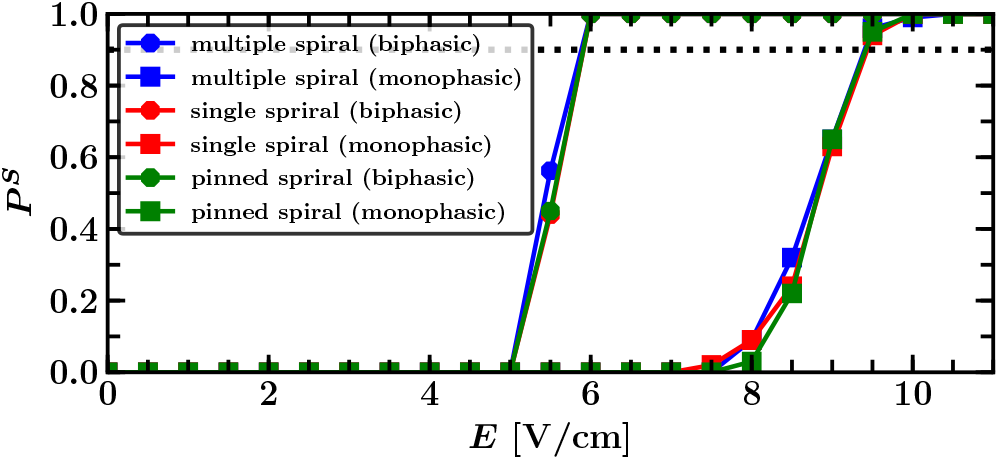
Single pulse termination probabilities *P* ^S^ for multiple spirals (blue), a single spiral (red) and a single pinned spiral (green) as functions of the applied field strength *E* and for biphasic (circles) and monophasic (squares) pulse shapes with 10 ms duration. The dotted horizontal line indicates a termination probability of 90% and defines *E*^90^, i.e., the minimum electric field strength to obtain a termination probability of at least 90%.

### B. Single pulse defibrillation

We determined the single pulse termination probability *P* ^S^, which is defined as the probability that a fibrillation state is converted into a quiescent state by a single defibrillation pulse, as a function of field strength *E*. We computed *P*^*S*^ as the fraction of successful termination events by running *N* = 50 simulations (see supplementary material) with different initial states for each parameter configuration, see Fig 3. The statistic error of *P* ^S^ is given by 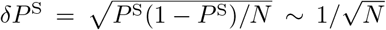. We have checked both monophasic and biphasic protocols, determining that biphasic (3 ms forward, 7 ms backward) are more effective than monophasic (10 ms forward) pulses, see Fig 3. This is in line with earlier findings^35^ and our previous results for different models exhibitng spatiotemporal chaos (=electrical turbulence)^27^. The probability to terminate a single free or pinned spiral is found to be almost identical than the one for terminating multiple spirals. Furthermore, the minimum electric field strength to reach a termination probability of at least 90% is nearly identical independent of the initial state. We found 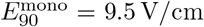 for monophasic and 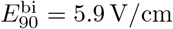 for biphasic pulses. Our results roughly compare with systematic experiments with monophasic pulses applied to over 200 fibrillation episodes of canine hearts *in vivo*. While defibrillation is hopeless for field strengths below 2.9 V/cm, the probability to defibrillate increases for higher values and is guaranteed for field strength above 8.5 V/cm.^32^

### C. Periodic anti-fibrillation pacing

We employed a sequence of biphasic pulses to mimick the experimental LEAP approach, since they require a lower field strength for defibrillation compared to single monophasic pulses, see Sec III B.

Taking a first look on successful LEAP of a single pinned spiral (Fig 4, we observe on the one hand that the spiral splits first up into smaller spirals until they get consecutively eliminated and on the other hand that the by each pulse excited tissue fraction increases with every pulse. Furthermore, we could not observe an unpinning process of the pinned spiral nor a shifting of the free spirals to the boundary. Similar observations could be made for multiple or a single free spiral, see provided videos in the supplementary material. This indicates already that neither the number of spirals nor the fact whether a spiral is pinned or not plays an essential role in the success of LEAP.

**FIG. 4.**
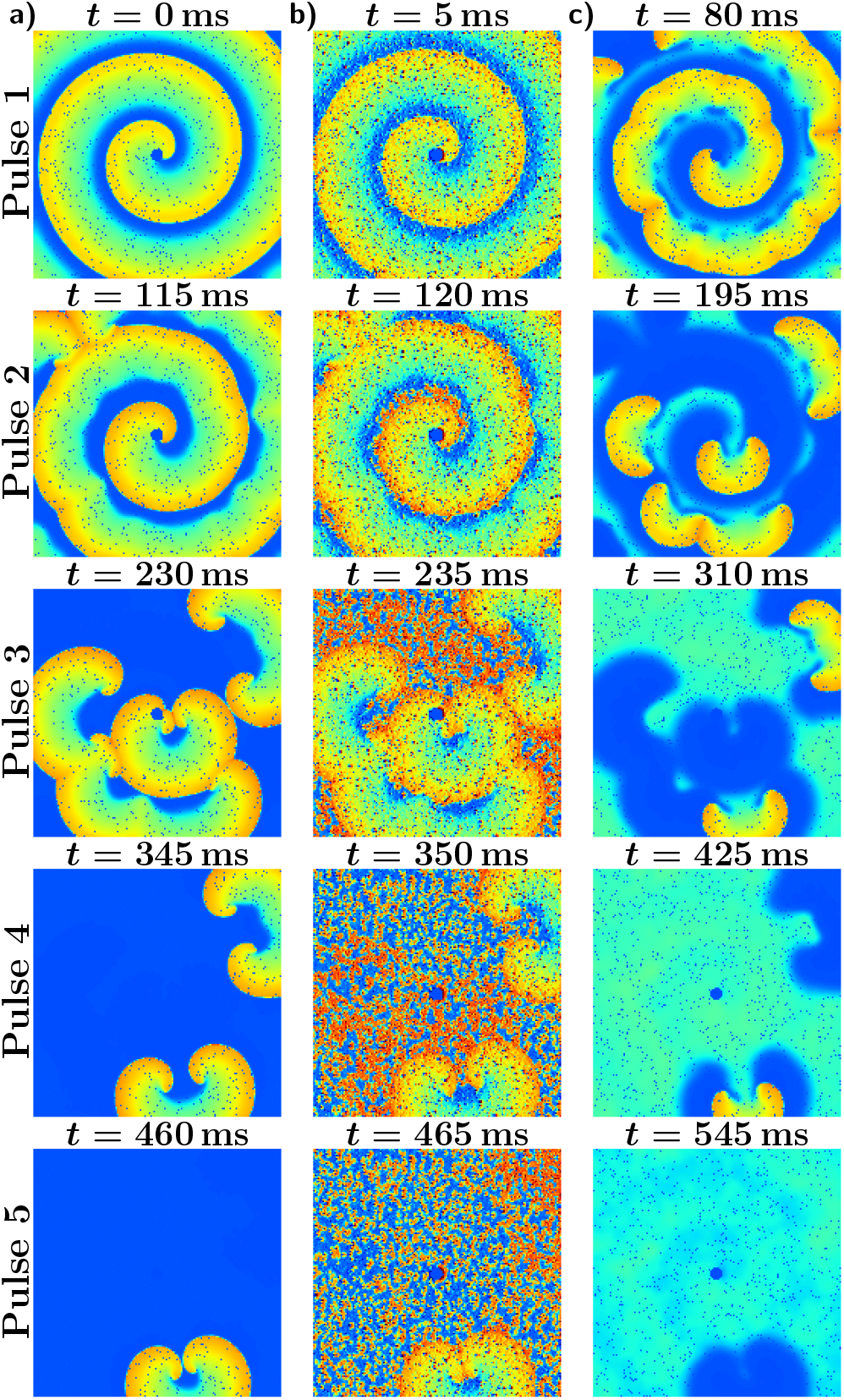
Sequence of snapshots for successful pacing with a period of *T/τ*_0_ = 1.15 and field strength *E* = 3.5 V/cm of a single pinned spiral, whose core is pinned to a circular heterogeneity at the center. The snapshots show the transmembrane potential right before, during, and 80 ms after the first, second, third, fourth and fifth pulse. (a) State right before a pulse. (b) The pulses induce excitation fronts at the big heterogeneities, which excite quickly nearly the entire excitable tissue, preventing the spirals to spread. (c) As a result of this, big spirals fall apart into smaller spirals while small spirals might get completely eliminated. Overall the by each pulse excited tissue fraction increases with every pulse until all excitation fronts are terminated.

The choice of the pacing period *T* turns out to be crucial for LEAP success, see Fig 5. The maximal termination probability *P* ^L^ is found for underdrive pacing with a period of *T*_max_ = 115 ms, which is 15% higher than the dominant period *τ*_0_ = 100 ms corresponding to the rotation period of stable spirals in the medium, see Fig 2. Further local maxima are found for *T*_max_/2 and 2 *T*_max_ albeit with considerable lower termination probabilities. The termination probability *P* ^L^ increased substantially with the number of pulses *n*. It is worth noting, that this picture is remarkable close to the success probabilities for defibrillation found in our earlier study using models exhibiting electrical turbulence^27^, though therein shape of the observed peaks was considerably more smeared out. Choosing the optimal pacing period *T*_max_, the minimal field strengths 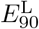 for a defibrillation success probability of at least 90% increases monotonously with the number of pulses *n*, see the colored dotted lines in Fig 6. We found for multiple spirals values of 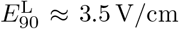 for five, 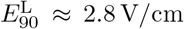 for ten and 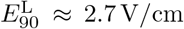 for fifteen pulses. Further substantial reductions for more pulses are unlikely as the difference between fifteen and ten pulses is already very small.

**FIG. 5.**
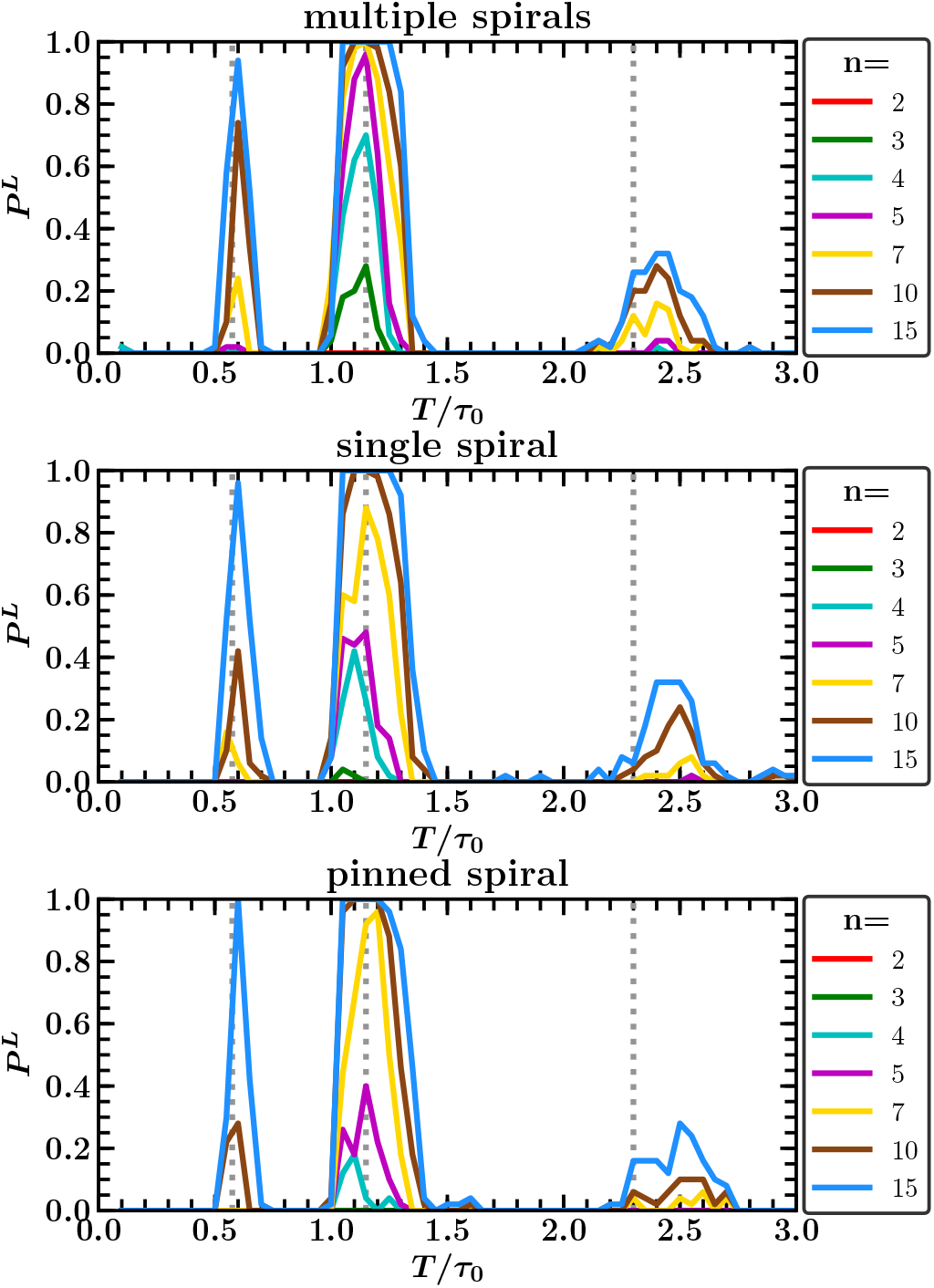
Termination probability *P* ^L^ of periodic pacing for multiple spirals, a single spiral and a single pinned spiral as a function of pacing period *T* for different pulse numbers *n* and fixed electric field strength *E* = 3.5 V/cm. Each curve exhibits an absolute maximum at *T*_max_ = 115 ms and pronounced local maxima near *T*_max_/2 and 2 *T*_max_ (vertical dotted lines). *T*_max_ = 115 ms is higher than the dominant period *τ*_0_ = 100 ms, indicating that underdrive pacing is optimal. Corresponding movies of simulations are provided in the supplementary material.

**FIG. 6.**
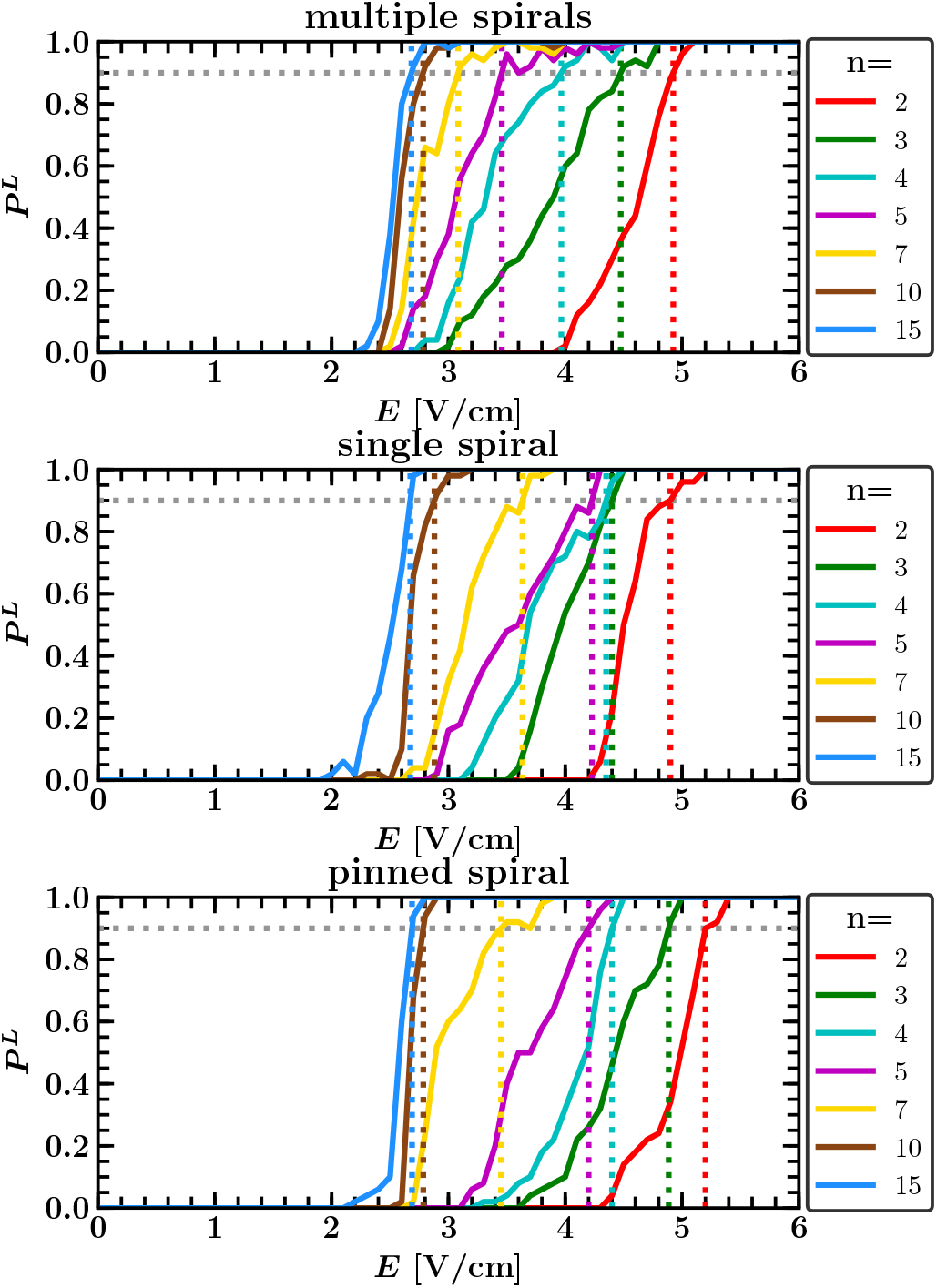
Termination probability *P* ^L^ of periodic pacing for multiple spirals, a single spiral and a single pinned spiral as a function of field strength *E* for different pulse numbers *n* and fixed pacing period *T* at *T*_max_ = 115 ms. The colored dotted lines indicate where the curves for different *n* first exceed 90% termination probability. For each configuration the probability to terminate multiple spirals is higher than to terminate a single free or pinned spiral. Corresponding movies of simulations are provided in the supplementary material.

The overall energy delivered by a periodic pacing protocol is proportional to the number of pulses *n* and the square of the electric field strength *E*. Compared to the field strength 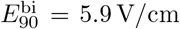 required for a single defibrillation pulse, the total required Energy for LEAP is always higher then for a single pulse. However there is a reduction of dissipated energy per pulse, which is for multiple spirals about 65% for five, 77% for ten and 79% for fifteen pulses.

The required energies for a single free or pinned stable spiral are for more then three pulses almost identical and only a little bit higher then the one required for multiple spirals. Hence, neither the number of spirals nor the fact whether a spiral is pinned or not seems to play an essential role for the success of periodic pacing.

### D. Analysis of periodic pacing dynamics

In the following we will only look at periodic pacing protocols with up to *n* = 5 pulses as the qualitative behavior does not change much for higher number of pulses. If not mentioned otherwise, averages and the termination probability *P*^*L*^ are computed by running *N* = 50 simulations with different initial states for each parameter configuration. Moreover we will only show the results for multiple spirals as the results for a single free or pinned spiral were similar.

## 1. Optimal pacing period

The range of successful pacing periods *T*, for which defibrillation is found in the simulations, becomes wider for high electric field strengths *E*, see Fig. 7a. We define the lower bound 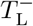 and upper bound 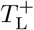 of this range as the two positions with 80% of the maximal possible termination probability:

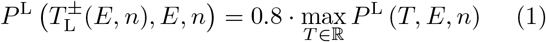

**FIG. 7.**
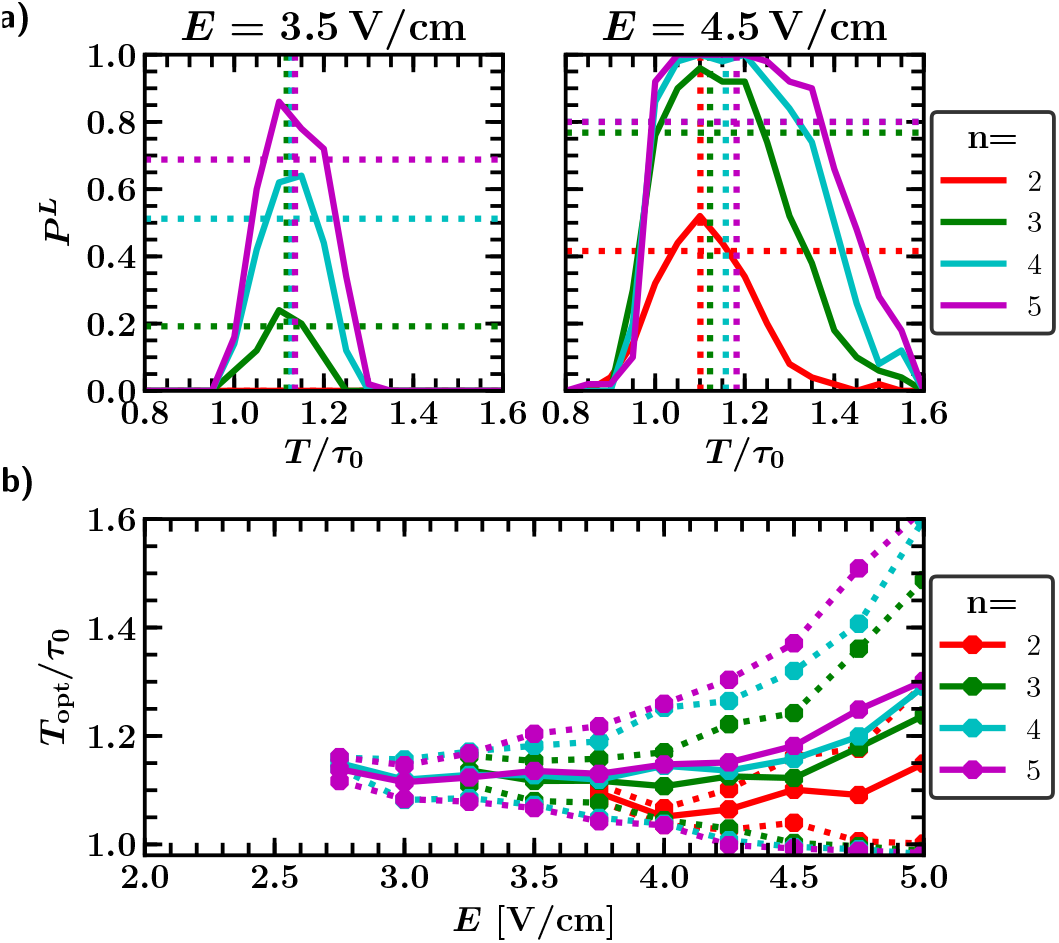
(a) Termination probability *P* ^L^ as a function of pacing period *T* and different pulse numbers *n* at fixed field strength *E* = 3.5 V/cm (left) and *E* = 4.5 V/cm (right). We define the lower and upper bound 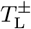of the range of successful pacing periods as the two positions with 80% of the maximal possible termination probability (intersection with the horizontal colored dotted lines). The vertical dotted lines denote the location of the optimal pacing period *T*_opt_, defined as the mean of those two bounds. (b) Optimal pacing period *T*_opt_ and the corresponding upper and lower bounds 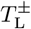 of the region of successful pacing periods as a function of electric field strength *E* and different pulse numbers *n*.

This allows us to study pacing protocols in regimes with low termination probabilities as well, compared to take the two positions at a fixed termination probability. Taking the global maximum *T*_max_ is not a good characterization for the optimal pacing period since it might not be located at the center of the peak. Therefore we define the optimal period *T*_*opt*_ as the mean of the two bounds 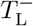 and 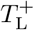:

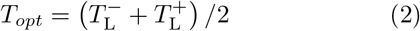

The range of pacing periods *T* with more than 80 % success rate starts at a minimal field strength, where the lower and upper bounds intersect, and widens monotonously with increasing field strength *E*, see Fig 7b. For low *E*, the lower and upper bounds both increase at roughly equal rate. However, for high *E* the lower bound remains almost constant, while the upper bound continuous to increase. As a result of that, the optimal period remains first constant and increases gradually for high *E*. For increasing number of pulses *n* the region of successful pacing periods becomes wider, while the minimal field strength decreases.

## 2. Prediction of the optimal pacing period

One can observe that during successful periodic pacing big spirals disintegrate into smaller spirals while small spirals get successively eliminated with each pulse until everything is terminated, see Fig 4. Hence, a progressively decrease of the total size of the spirals, given by the complement of the fraction of excitable tissue *F*_Exc_ seems to be the crucial factor for the success of periodic pacing. Successful pacing protocols are characterized by high fractions of excitable tissue *F*_Exc_ right before the pulses.

Achieving a maximal fraction of excitable tissue *F*_Exc_ right before a pulse is a trade-off between pacing fast enough in order to prevent already recovered tissue to get excited by remaining excitation fronts and pacing slow enough, giving the tissue enough time to fully recover. As shown in Fig 8a, the range of pacing periods *T* with a high average fraction of excitable tissue right before a pulse ⟨*F*_Exc_⟩ becomes for an increasing pulse number *n* and electric field strength *E* wider towards higher values of *T*. That is, since the total size of the remaining excitation waves decreases during successful pacing, see Fig 4a, slowing down the excitation of excitable tissue. This widening of the range of pacing periods *T* with a high average fraction of excitable tissue right before a pulse ⟨*F*_Exc_⟩ towards higher values of *T* looks similar to the one for the range of high termination probabilities *P* ^L^, see Fig 7a. As shown in Fig 8b, the average fraction of excitable tissue right before a pulse ⟨*F*_Exc_⟩ is highly correlated with the termination probability *P* ^L^ of a pulse, at least for pulse numbers *n* ≥ 3. A good estimation of the optimal pacing period *T*_opt_, which is optimal in the sense of a high termination probability *P* ^L^, should be therefore given by a optimal pacing period with respect to high ⟨*F*_Exc_⟩. We define this pacing period 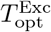 analogous to *T*_opt_ as the mean of the lower and upper bound 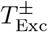 of pacing periods *T* achieving a ⟨*F*_Exc_⟩ of at least 80% of the maximal possible (*F*_Exc_):

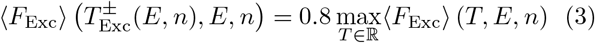

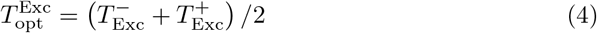

**FIG. 8.**
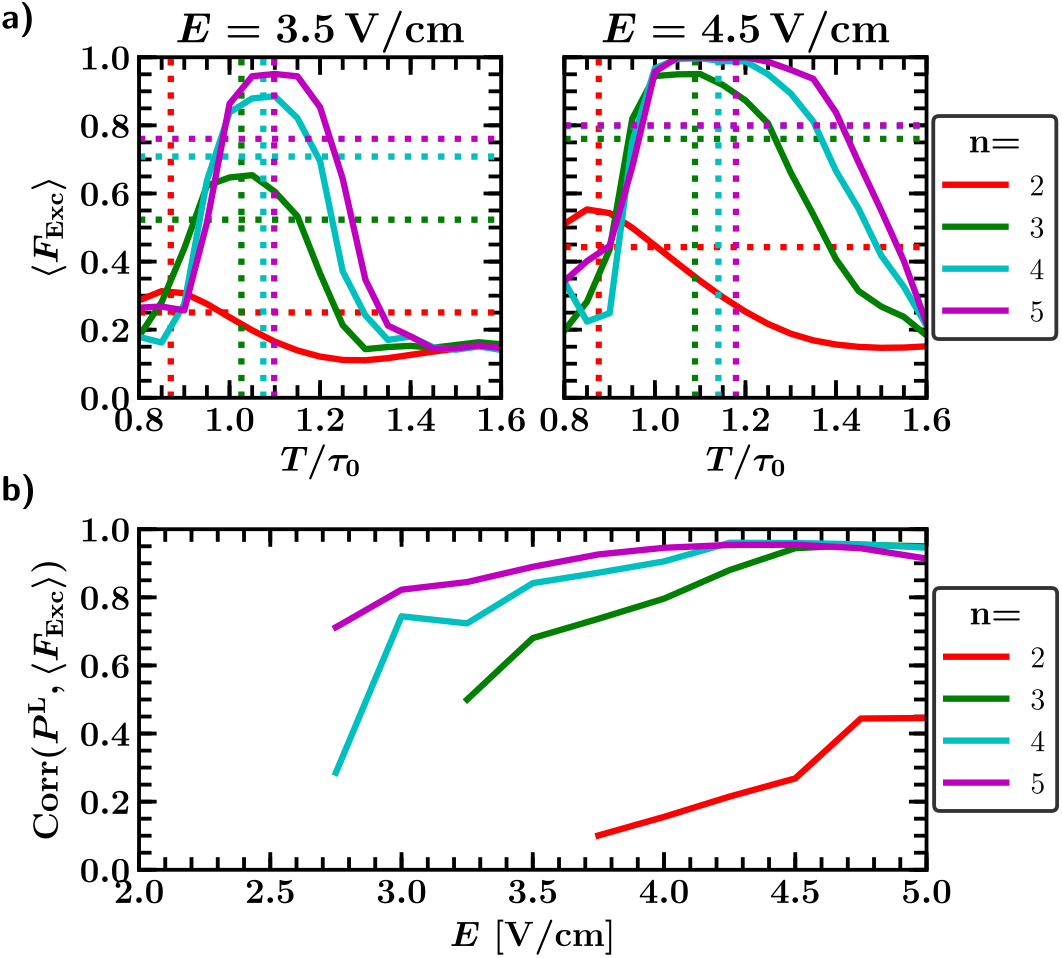
Average fraction of excitable tissue right before a pulse ⟨*F*_Exc_⟩ (a) as a function of pacing period *T* and pulse number *n* for a LEAP protocol with fixed field strength *E* = 3.5 V/cm (left) and *E* = 4.5 V/cm (right). The vertical dotted lines in denote the location of the optimal pacing period 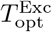 with respect to a high fraction of excitable tissue right before a pulse *F*_Exc_, defined as the mean of the two positions with 80% of the maximal possible value of ⟨*F*_Exc_⟩ (intersection with the horizontal colored dotted lines). (b) Pearson correlation coefficient between the termination probability *P* ^L^, see Fig 7a, and the average excitable tissue fraction ⟨*F*_Exc_⟩ with respect to the pacing period *T* as a function of field strength *E* and pulse number *n*. Both quantities are relatively high correlated for pulse numbers *n ≥* 3.

As shown in Fig 9a, 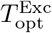 is a good estimation of the optimal pacing period *T*_opt_ and coincides for pulse numbers *n* ≥ 5 almost completely with *T*_opt_. This also applies for lower *n* in the regime of high field strength *E* and 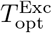 lies for this field strength *E*, where the correlation coefficient of *P*_L_ and ⟨*F*_Exc_⟩ is greater then 0.8, see Fig 8c, still in the regime of successful pacing periods.

**FIG. 9.**
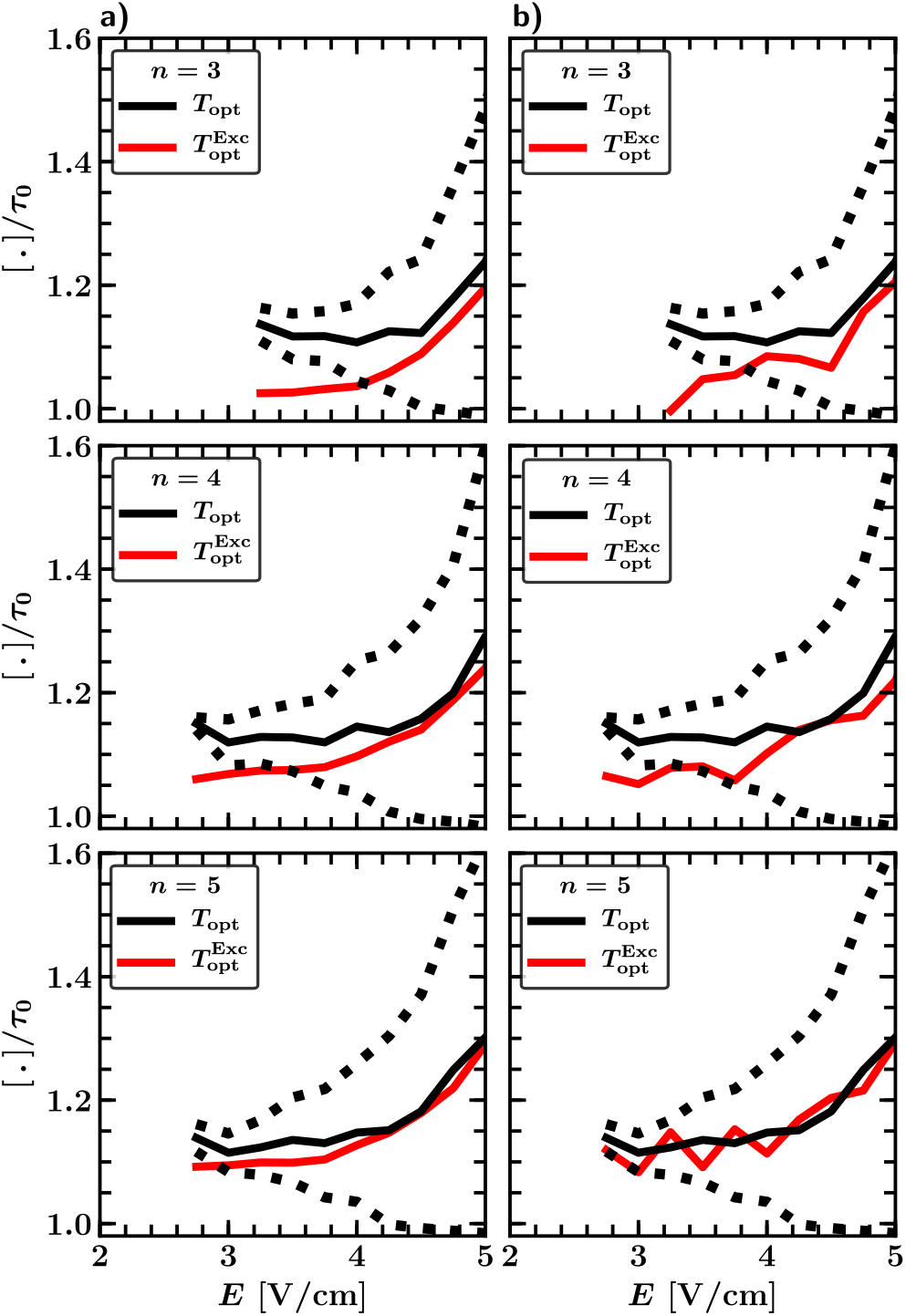
(a) Optimal pacing period *T*_opt_ (black) with respect to a high termination probability *P* ^L^ and optimal pacing period *T* ^Exc^ (red) with respect to a high average of the by pulse excited tissue fraction ⟨*F*_Exc_⟩ as a function of field strength *E* and different pulse numbers *n.* 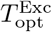 is a good estimation of *T*_opt_ for high *E*. It lies for *n* = 3 and *n* = 4 and field strength lower then 4 V/cm and 3.4 V/cm respectively outside the regime of successful pacing periods, however, coincides for *n ≥* 5 almost completely with *T*_opt_. (b) Same as in (a), however, ⟨*F*_Exc_⟩ is here determined by only one randomly chosen initial condition and not as an average for every parameter configuration (*E, T, n*). Nevertheless, the impact on 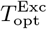 are not significant and 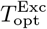 keeps a nearly equally good estimation of *T*_opt_ compared to the one from (a) for an averaged ⟨*F*_Exc_⟩.

Measuring the fraction of excitable tissue right before a pulse *F*_Exc_ is harder then to determine whether a pulse is successful or not, especially in a real experiment. Therefore, it would not be worth to determine the optimal period *T*_opt_ by the use of ⟨*F*_Exc_⟩ instead of the success probability *P* ^L^. However the fraction of excitable tissue right before a pulse *F*_Exc_ of a single pulse can be measured directly, while *P* ^L^ can be only determined as the mean of successful (0) and unsuccessful (1) events. This might allow us to determine an accurate value for *F*_Exc_ with less simulations or experiments and determining *F*_Exc_ by only one randomly chosen initial condition for every parameter configuration has already no significant impact on 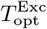, as shown in Fig 9b. 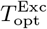 still coincides for pulse numbers *n* ≥ 5 quiet good with *T*_opt_ and is only a little bit noisy.

## IV. CONCLUSION AND DISCUSSION

We have carried out a statistical study of simulations of periodic pacing for an electrophysiological model exhibiting stable spirals within a two-dimensional model of cardiac tissue perforated by blood vessels. It turns out that underdrive pacing (= pacing with a period lower than the spiral rotation period) results in maximal energy reduction and that neither the number of spirals nor the fact whether a spiral is pinned or not plays an essential role in the defibrillation success of periodic pacing. A protocol with ten pulses and a pacing period 15% higher than the dominant period requires 77% less energy per pulse than a single biphasic shock for successful defibrillation. We observed that successful pacing protocols are characterized by repeated pulsing at times when the fraction of excitable tissue is high. This fraction of excitable tissue right before a pulse is found to be highly correlated with the termination probability allowing an efficient alternative to estimate the optimal pacing period to achieve defibrillation. The appropriate choice of length scales is an important issue. We have chosen a scaling based on the experimental observation regarding the sizes of the heterogeneities in^8^ and picked a value of the conductivity such that the single pulse defibrillation threshold reproduces experimentally found values^8,32^. Furthermore, the size of the system was picked sufficiently large to generate initial states not only with a single free or pinned spiral but also with around ten coexisting spirals. While the initial dynamics after the start of pacing depended strongly on the initial conditions the success probability after a sufficiently large number of pulses was similar for all chosen initial conditions.

We found that defibrillation by periodic pacing works most efficiently with underdrive pacing for stable spirals compared to our results for spatiotemporal chaos^27^ where resonant pacing was favorable at low field strength and underdrive pacing was only superior for stronger electrical field. An increase of the optimal pacing period with the field strength is also observable for stable spirals. This is caused by an expansion and not by a shift of the regime of successful pacing towards higher values of the period *T* and is in line to our earlier observation for defibrillation by periodic pacing in models exhibits spatiotemporal chaotic electric activity^27^. A protocol with 10 pulses and a pacing period 15% higher than the dominant period requires 77% less energy per pulse than a single biphasic shock for successful defibrillation, which corresponds with the energy reductions between 75% and 90%, found for successful underdrive pacing of canine hearts^8^.

Current theoretical approaches to understand LEAP for stable spirals usually focus on the process of unpinning and the removal of free stable spirals by shifting the spiral tips to the boundary of the heart.^15,16,20,21^ These theories predict also optimal success for underdrive pacing with periods between 110% and 120%,^23,24^. However, our observation, that neither the initial number of spirals nor the fact whether a spiral is pinned or not plays an essential role in the success of periodic pacing, see Fig 5 and 6 point towards different essential ingredients. Furthermore we found in our simulations that the process of unpinning and shifting of the spirals to the boundary of the heart did not play an essential role, see Fig 4 and provided videos in the supplementary material. In contrast, we observed that during successful pacing big spirals disintegrate into smaller spirals while small spirals get successively eliminated with each pulse until all activity is terminated, see Fig 4.

Therefore, a progressive decrease of the total size of the spirals in parallel to an increase of the fraction of excitable tissue *F*_Exc_ turned out to be the crucial factor for the success of periodic pacing. In other words, successful pacing protocols are characterized by pulsing at times when the fraction of excitable tissue *F*_Exc_ is high. This picture is consistent with observations, made in experimental^8,25^ and numerical studies^25,26^, which interpret repeat pacing as a synchronization process of the phases of the electrical activity in the tissue. The subsequent increase in the total fraction of excitable tissue with each pulse in the periodic pacing leads to an increasing synchronisation of local excitation cycles, which is, however, depending on the simultaneous trigger provided by the pacing and therefore different from the more frequently studied mutual synchronisation occuring in oscillatory media.

The observation that the termination probability *P*_L_ of a pulse is highly correlated with the fraction of excitable tissue right before a pulse *F*_Exc_ for pulse numbers *n* ≥ 3, see Fig 8b allows us to determine a criterion for the optimal pacing period *T*_opt_ (optimality here refers a high termination probability *P* ^L^). This done by finding the pacing period that maximizes *F*_Exc_, see Fig 9a. The big advantage of this estimation is the fact, that the excitable fraction *F*_Exc_ is easily accessible and could be measured directly in experiments, whereas the termination probability *P* ^L^ can be only determined by counting successful (0) and unsuccessful (1) events requiring therefore a large amount of independent measurements. In our context, determining *F*_Exc_ by only one simulation for every parameter configuration provides a very good approximation of the optimal pacing period, see Fig 9b. Finally, it is worth pointing out that in this study of an electrophysiological model that represents an excitable medium wherein one or more stable spirals can coexist in the absence of pacing is in many ways very similar to the results that we found earlier for two electrophysiological models exhibiting spatiotemporal chaos (= electrical turbulence)^27^. Therefore, the mechanistic considerations described in the last part of the results section above may generalize to these qualitatively different dynamics.

## SUPPLEMENTARY MATERIAL

See supplementary material for movies of the simulations. The simulation domain has in all movies an edge length of about ten times the correlation length *ξ* and no-flux boundary condition. In movies 3-8, the (first) pulse is applied at *t* = 0 ms. All biphasic pulses are applied for 7 ms in the forward direction and 3 ms in the backward direction.

Movie 1: State multiple spirals Movie 2: State of a single spiral

Movie 3: State of a single pinned spiral

Movie 4: Defibrillation of multiple spirals with a single biphasic pulse for *E* = 6 V/cm

Movie 5: Defibrillation of a single spiral with a single biphasic pulse for *E* = 6 V/cm

Movie 6: Defibrillation of a single pinned spiral with a single biphasic pulse for *E* = 6 V/cm

Movie 7: Defibrillation of multiple spirals by LEAP with five biphasic pulses for *E* = 3.5 V/cm and pacing period *T* = 115 ms

Movie 8: Defibrillation of a single spiral by LEAP with five biphasic pulses for *E* = 3.5 V/cm and pacing period *T* = 115 ms

Movie 9: Defibrillation of a single pinned spiral by LEAP with five biphasic pulses for *E* = 3.5 V/cm and pacing period *T* = 115 ms

## ACKNOWLEDGMENTS

We acknowledge financial support by DFG through funding of project B5 in SFB 910 and acknowledge useful discussions with Harald Engel, Valentin Krinsky, Stefan Luther and Ulrich Parlitz.

## References

1 R. A. Gray, A. M. Pertsov, and J. Jalife, “Spatial and temporal organization during cardiac fibrillation,” Nature 392, 75–78 (1998).

2 F. X. Witkowski, L. J. Leon, P. A. Penkoske, W. R. Giles, M. L. Spano, W. L. Ditto, and A. T. Winfree, “Spatiotemporal evolution of ventricular fibrillation,” Nature 392, 78–82 (1998).

3 M. P. Nash, A. Mourad, R. H. Clayton, P. M. Sutton, C. P. Bradley, M. Hayward, D. J. Paterson, and P. Taggart, “Evidence for multiple mechanisms in human ventricular fibrillation,” Circulation 114, 536–542 (2006).

4 K. H. Ten Tusscher and A. V. Panfilov, “Organization of ventricular fibrillation in the human heart,” Circulation Research 100, e87–e101 (2007).

5 W. H. Maisel, “Pacemaker and icd generator reliability: metaanalysis of device registries,” Jama 295, 1929–1934 (2006).

6 G. P. Walcott, C. R. Killingsworth, and R. E. Ideker, “Do clinically relevant transthoracic defibrillation energies cause myocardial damage and dysfunction?” Resuscitation 59, 59–70 (2003).

7 F. H. Fenton, S. Luther, E. M. Cherry, N. F. Otani, V. Krinsky, A. Pumir, E. Bodenschatz, and R. F. Gilmour, “Termination of atrial fibrillation using pulsed low-energy far-field stimulation,” Circulation 120, 467–476 (2009).

8 S. Luther, F. H. Fenton, B. G. Kornreich, A. Squires, P. Bittihn, D. Hornung, M. Zabel, J. Flanders, A. Gladuli, L. Campoy, E. M. Cherry, G. Luther, G. Hasenfuss, V. I. Krinsky, A. Pumir, R. F. G. Jr, and E. Bodenschatz, “Low-energy control of electrical turbulence in the heart,” Nature 475, 235–239 (2011).

9 C. M. Ambrosi, C. M. Ripplinger, I. R. Efimov, and V. V. Fedorov, “Termination of sustained atrial flutter and fibrillation using low-voltage multiple-shock therapy,” Heart Rhythm 8, 101–108 (2011).

10 W. Li, A. H. Janardhan, V. V. Fedorov, Q. Sha, R. B. Schuessler, and I. R. Efimov, “Low-energy multistage atrial defibrillation therapy terminates atrial fibrillation with less energy than a single shockclinical perspective,” Circulation: Arrhythmia and Electrophysiology 4, 917–925 (2011).

11 W. Li, C. M. Ripplinger, Q. Lou, and I. R. Efimov, “Multiple monophasic shocks improve electrotherapy of ventricular tachycardia in a rabbit model of chronic infarction,” Heart Rhythm 6, 1020–1027 (2009).

12 A. H. Janardhan, W. Li, V. V. Fedorov, M. Yeung, M. J. Wallendorf, R. B. Schuessler, and I. R. Efimov, “A novel low-energy electrotherapy that terminates ventricular tachycardia with lower energy than a biphasic shock when antitachycardia pacing fails,” Journal of the American College of Cardiology 60, 2393–2398 (2012).

13 S. Weidmann, “Effect of current flow on the membrane potential of cardiac muscle,” The Journal of Physiology 115, 227 – 236 (1951).

14 J. P. Wikswo Jr, S.-F. Lin, and R. A. Abbas, “Virtual electrodes in cardiac tissue: a common mechanism for anodal and cathodal stimulation.” Biophysical Journal 69, 2195 (1995).

15 A. Pumir and V. Krinsky, “Unpinning of a rotating wave in cardiac muscle by an electric field,” Journal of theoretical biology 199, 311–319 (1999).

16 A. Pumir, V. Nikolski, M. Hörning, A. Isomura, K. Agladze, K. Yoshikawa, R. Gilmour, E. Bodenschatz, and V. Krinsky, “Wave emission from heterogeneities opens a way to controlling chaos in the heart,” Physical review letters 99, 208101 (2007).

17 P. Bittihn, M. Hörning, and S. Luther, “Negative curvature boundaries as wave emitting sites for the control of biological excitable media,” Physical review letters 109, 118106 (2012).

18 B. J. Caldwell, M. L. Trew, and A. M. Pertsov, “Cardiac response to low energy field pacing challenges the standard theory of defibrillation,” Circulation: Arrhythmia and Electrophysiology, CIRCEP–114 (2015).

19 J. Christoph, M. Chebbok, C. Richter, J. Schröder-Schetelig, P. Bittihn, S. Stein, I. Uzelac, F. Fenton, G. Hasenfuß, R. Gilmour Jr, et al., “Electromechanical vortex filaments during cardiac fibrillation,” Nature 555, 667 (2018).

20 S. Takagi, A. Pumir, D. Pazo, I. Efimov, V. Nikolski, and V. Krinsky, “Unpinning and removal of a rotating wave in cardiac muscle,” Physical review letters 93, 058101 (2004).

21 C. M. Ripplinger, V. I. Krinsky, V. P. Nikolski, and I. R. Efimov, “Mechanisms of unpinning and termination of ventricular tachycardia,” American Journal of Physiology-Heart and Circulatory Physiology 291, H184–H192 (2006).

22 A. Pumir, S. Sinha, S. Sridhar, M. Argentina, M. Hörning, S. Filippi, C. Cherubini, S. Luther, and V. Krinsky, “Wave-train-induced termination of weakly anchored vortices in excitable media,” Physical Review E 81, 010901 (2010).

23 T. Shajahan, S. Berg, S. Luther, V. Krinski, and P. Bittihn, “Scanning and resetting the phase of a pinned spiral wave using periodic far field pulses,” New Journal of Physics 18, 043012 (2016).

24 D. Hornung, V. Biktashev, N. Otani, T. Shajahan, T. Baig, S. Berg, S. Han, V. Krinsky, and S. Luther, “Mechanisms of vortices termination in the cardiac muscle,” Open Science 4, 170024 (2017).

25 Y. C. Ji, I. Uzelac, N. Otani, S. Luther, R. F. Gilmour Jr, E. M. Cherry, and F. H. Fenton, “Synchronization as a mechanism for low-energy anti-fibrillation pacing,” Heart rhythm 14, 1254–1262 (2017).

26 L. J. Rantner, B. M. Tice, and N. A. Trayanova, “Terminating ventricular tachyarrhythmias using far-field low-voltage stimuli: mechanisms and delivery protocols,” Heart Rhythm 10, 1209–1217 (2013).

27 P. Buran, M. Bär, S. Alonso, and T. Niedermayer, “Control of electrical turbulence by periodic excitation of cardiac tissue,” Chaos: An Interdisciplinary Journal of Nonlinear Science 27, 113110 (2017).

28 L. Luo and Y. Rudy, “A model of the ventricular cardiac action potential: depolarization, repolarization and their interaction,” Circ. Res. 68, 1501–1526 (1991).

29 F. Fenton and A. Karma, “Vortex dynamics in three-dimensional continuous myocardium with fiber rotation: filament instability and fibrillation,” Chaos: An Interdisciplinary Journal of Nonlinear Science 8, 20–47 (1998).

30 F. Xie, Z. Qu, A. Garfinkel, and J. N. Weiss, “Electrophysiological heterogeneity and stability of reentry in simulated cardiac tissue,” American Journal of Physiology-Heart and Circulatory Physiology 280, H535–H545 (2001).

31 J. P. Keener and J. Sneyd, Mathematical physiology, vol. 1 (Springer, 2009).

32 F. X. Witkowski, P. A. Penkoske, and R. Plonsey, “Mechanism of cardiac defibrillation in open-chest dogs with unipolar dc-coupled simultaneous activation and shock potential recordings.” Circulation 82, 244–260 (1990).

33 I. R. Efimov, M. W. Kroll, and P. Tchou, Cardiac bioelectric therapy: mechanisms and practical implications (Springer Science & Business Media, 2008).

34 A. Winfree, “Evolving perspectives during 12 years of electrical turbulence,” Chaos: An Interdisciplinary Journal of Nonlinear Science 8, 1–19 (1998).

35 J. Bragard, A. Simic, J. Elorza, R. O. Grigoriev, E. M. Cherry, R. F. Gilmour Jr, N. F. Otani, and F. H. Fenton, “Shock-induced termination of reentrant cardiac arrhythmias: Comparing monophasic and biphasic shock protocols,” Chaos: An Interdisciplinary Journal of Nonlinear Science 23, 043119 (2013).

